# Optimal methods for analyzing targeted pairwise knockout screens

**DOI:** 10.1101/2024.08.19.608665

**Authors:** Juihsuan Chou, Nazanin Esmaeili Anvar, Reem Elghaish, Junjie Chen, Traver Hart

## Abstract

**Background:** Synthetic lethality offers a promising strategy for cancer treatment by targeting genetic vulnerabilities unique to tumor cells, leading to selective tumor cell death. However, single-gene knockout screens often miss functional redundancy due to paralog genes. Multiplex CRISPR systems, including various Cas9 and Cas12a platforms, have been developed to assay genetic interactions, yet no systematic comparison of method to identify synthetic lethality from CRISPR screens has been conducted.

**Results:** We evaluated data from four in4mer CRISPR/Cas12a screens in cancer cell lines, using three bioinformatic approaches to identify synthetic lethal interactions: delta log fold change (dLFC), Z-transformed dLFC (ZdLFC), and rescaled dLFC (RdLFC). Both ZdLFC and RdLFC provided more consistent identification of synthetic lethal pairs across cell lines compared to the unscaled dLFC method.

**Conclusions:** The ZdLFC method offers a robust framework for scoring synthetic lethal interactions from paralog screens, providing consistent results across different cell lines without requiring a training set of known positive interactors.

## Introduction

Synthetic lethality offers an attractive approach to cancer treatment. Conceptually, mutations arising in tumors may give rise to genetic vulnerabilities that, when treated with targeted agents, result in tumor cell death with minimal effect on normal tissues. CRISPR-mediated genetic screens in over a thousand cell lines ^1–4^ have identified context-specific essential genes – candidate tumor-specific drug targets – but these single-gene knockout screens systematically miss functional buffering by paralogs ^5,6^.

To systematically understand paralog synthetic lethality, several groups have developed multiplex perturbation systems to assay genetic interactions in human cells. The various CRISPR platforms include dual Cas9, hybrid Cas9 and Cas12a, Cas12a-only, and orthologous Cas9 from S. aureus and S. pyogenes ^6–10^. Each study utilizes conceptually similar approaches to identify genetic interactions, by comparing single-gene knockout phenotypes to paired knockout phenotype. Despite this overall similarity in experimental design, each group employs different hit-calling pipelines. Thompson et al.^8^, Parrish et al.^7^, Dede et al.^6^, and Gonatopoulos-Pournatzis et al.^10^ quantify genetic interaction effects by calculating the delta log fold change (dLFC), defined as the difference between observed and expected log2 fold change (LFC). The expected LFC for paired gRNA constructs is calculated by summing the observed LFC values for individual gRNAs paired with non-targeting controls^7,8^, intergenic controls^10^, or nonessential controls^6^, depending on the library design. In Thompson et al.^8^, variance smoothing is then performed, and hits are identified using both t-tests and the robust ranking algorithm (RRA). Parrish et al.^7^ established a linear regression of control expected versus observed LFC and calculated the genetic interaction (GI) score as the residual of each observed LFC from the control regression line. Hits were identified by applying statistical significance tests and false discovery rate (FDR) correction. Dede et al.^6^ converted dLFC scores to Z-scores by truncating the top and bottom 2.5% of dLFC scores and identified hits with Z-transformed dLFC scores less than −3. Gonatopoulos-Pournatzis et al. used the Wilcoxon rank-sum test followed by Benjamini-Hochberg FDR correction to compare the observed LFC set to the expected LFC set for each gene pair. Ito et al. employed GEMINI^11^, a variational Bayesian method, to score GI. While these methods for analyzing multiplex CRISPR screens have successfully identified robust interactions that withstand subsequent validation, they are often tailored to specific library formats, limiting their generalizability across different experimental designs.

The in4mer platform is a CRISPR/Cas12a system for combinatorial gene knockout^12^. The key element of the in4mer system is a four-guide array of Cas12a guide RNA, expressed from a U6 promoter, that the Cas12a endonuclease can process and use as individually targeting gRNA, potentially resulting in multiple gene knockouts in the same cell. Using the in4mer platform, libraries targeting more than 2,000 paralog pairs were screened for synthetic lethality. Here we use data from in4mer screens in four cancer cell lines to evaluate different bioinformatic approaches for hit calling in synthetic lethal (SL) screens.

## Results and discussion

Synthetic lethality is a limiting case of genetic interaction, where joint perturbation phenotype is more severe than expected from the individual pairwise knockouts (Figure 1A). Paralogs represent a fruitful search space in which to test genetic interaction technologies, both experimental and informatic, because functional buffering by paralogs is far more frequent than genetic interactions between non-paralogous genes. However, there is no clear standard for analyzing this type of data. Here we consider the most straightforward approach, delta log fold change (dLFC), and two derivatives of this approach, a Z-transformed ZdLFC (Figure 1B) and a supervised, rescaled RdLFC (Figure 1C). Notably, the Z-score approach uses the observed distribution of dLFC to estimate a null model, and scores deviations from the null (high |Z| scores) as hits. The RdLFC uses the same null model and adds an empirical model for hits, based on the observed dLFC of the positive control reference set of 13 paralog synthetic lethals defined in Esmaeili Anvar et al^12^, and scales other gene pairs accordingly. We also considered a fourth approach, ParaBagel, which derives a Bayes Factor for synthetic lethality from these two models -- analogous to the Bagel algorithm^13,14^ for classifying essential genes from CRISPR knockout screens – but we found this approach unsuitable for the data (Supplementary method). We apply these three approaches to data from two Inzolia library screens in cancer cell lines, and two additional screens with an earlier version of the library, “Prototype” (Figure 1D; Esmaeili Anvar et al^12^), to evaluate hit-calling consistency across pipelines. A complete set of scores for all pairs is available in Supplementary Table 1.

**Figure 1.**
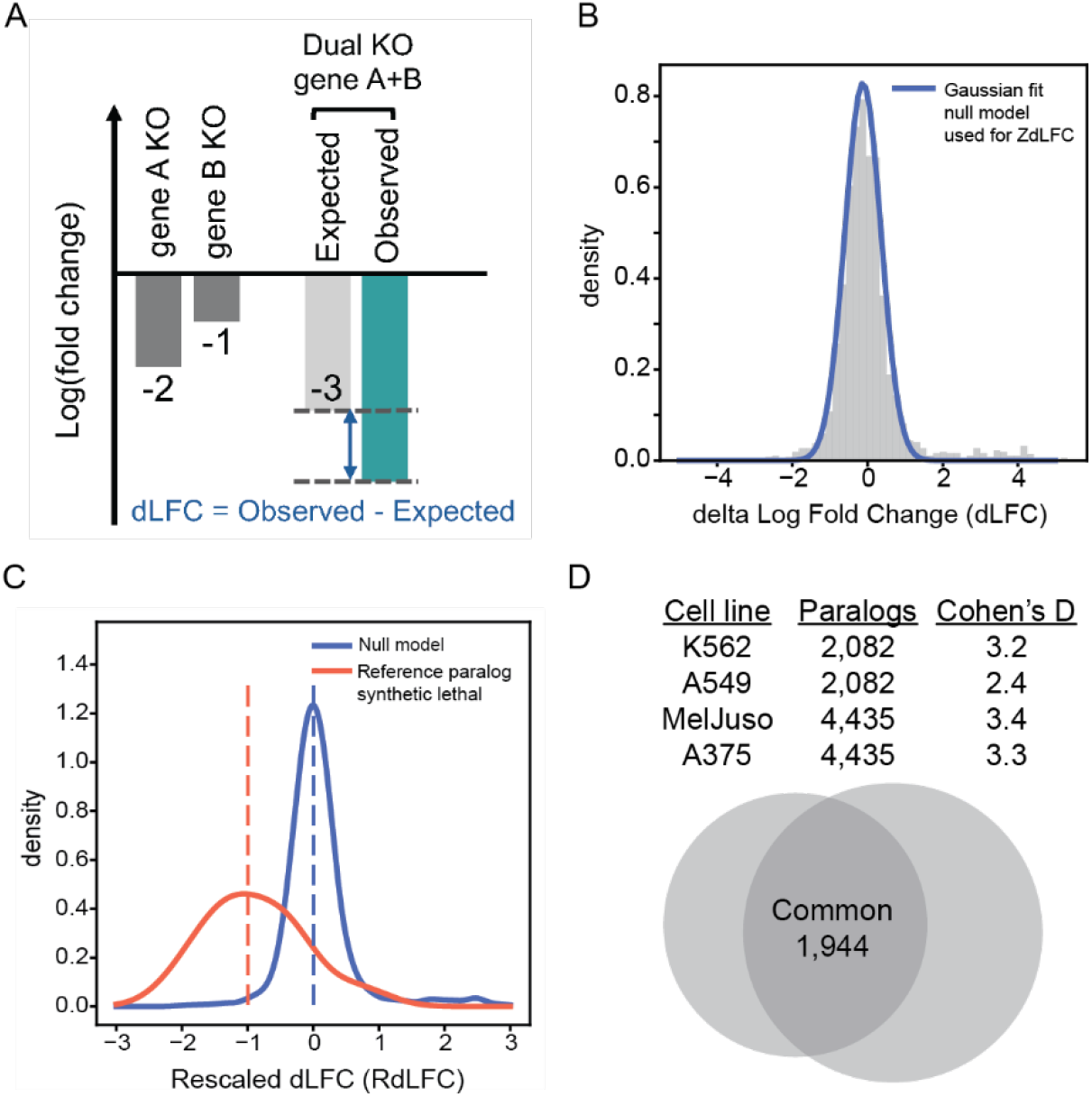
(A) Measuring synthetic lethality between paralog pairs. Single gene knockout (KO) fitness is determined by calculating the mean log fold change (LFC) of gRNAs targeting the specific gene. The expected dual gene KO fitness is the sum of the single gene KO LFCs for gene A and gene B. The Delta log fold change (dLFC) represents the difference between the observed and expected dual KO LFC. (B) The dLFC histogram of an Inzolia screen is shown with a normal distribution fit after removing outliers (Methods). The blue curve represents the fit of the null model, which is used to calculate the ZdLFC scores. (C) The dLFC distribution of an Inzolia screen after rescaling (Methods). The red and blue curves indicate kernel density plots of the 13 reference paralog synthetic lethal (positive control) and the null model (negative control), respectively. The dotted lines indicate that the median of positive controls is rescaled to −1, while the negative controls are set to 0. (D) Table displaying the four screens conducted with the Inzolia library: the “prototype” library in K562 and A549 cell lines, and the final Inzolia library in MelJuso and A375 cell lines. The table includes the number of paralogs in each screen and the Cohen’s D quality score, which measures the LFC differences between essential and nonessential controls relative to variability (Methods). The Venn diagram illustrates the number of common paralog pairs between the “prototype” and the final Inzolia library.

We applied these three methods to the 1,944 paralog pairs that were common to the two in4mer libraries, across the four cell lines screened. As in Esmaeili Anvar et al^12^, we reasoned that most paralog synthetic lethality would be common across various backgrounds, and we therefore sought to identify the computational approach that maximized this commonality. Using the dLFC approach, with a simple threshold of dLFC < −1 (strong genetic interaction) and LFC of the pair < −1 (pairwise knockout shows strong fitness defect), we identified 16 to 75 paralogs SL in the four cell lines (Figure 2A), with considerable overlap (Figure 2B). For ZdLFC, we defined a hit as ZdLFC < −2 and Z-transformed observed LFC (ZLFC, method) < −2 (Figure 2C, 2D). Finally, for RdLFC, we defined a hit as interaction score < −0.7 and same ZLFC < −2, with the interaction score threshold being somewhat arbitrarily chosen to yield roughly the same number of hits as the ZdLFC approach in order to minimize sample size as a source of bias when comparing the methods (Figure 2E, 2F). A visualization of the three methods as applied to the four screens is available in Supplementary Figure 1.

**Figure 2.**
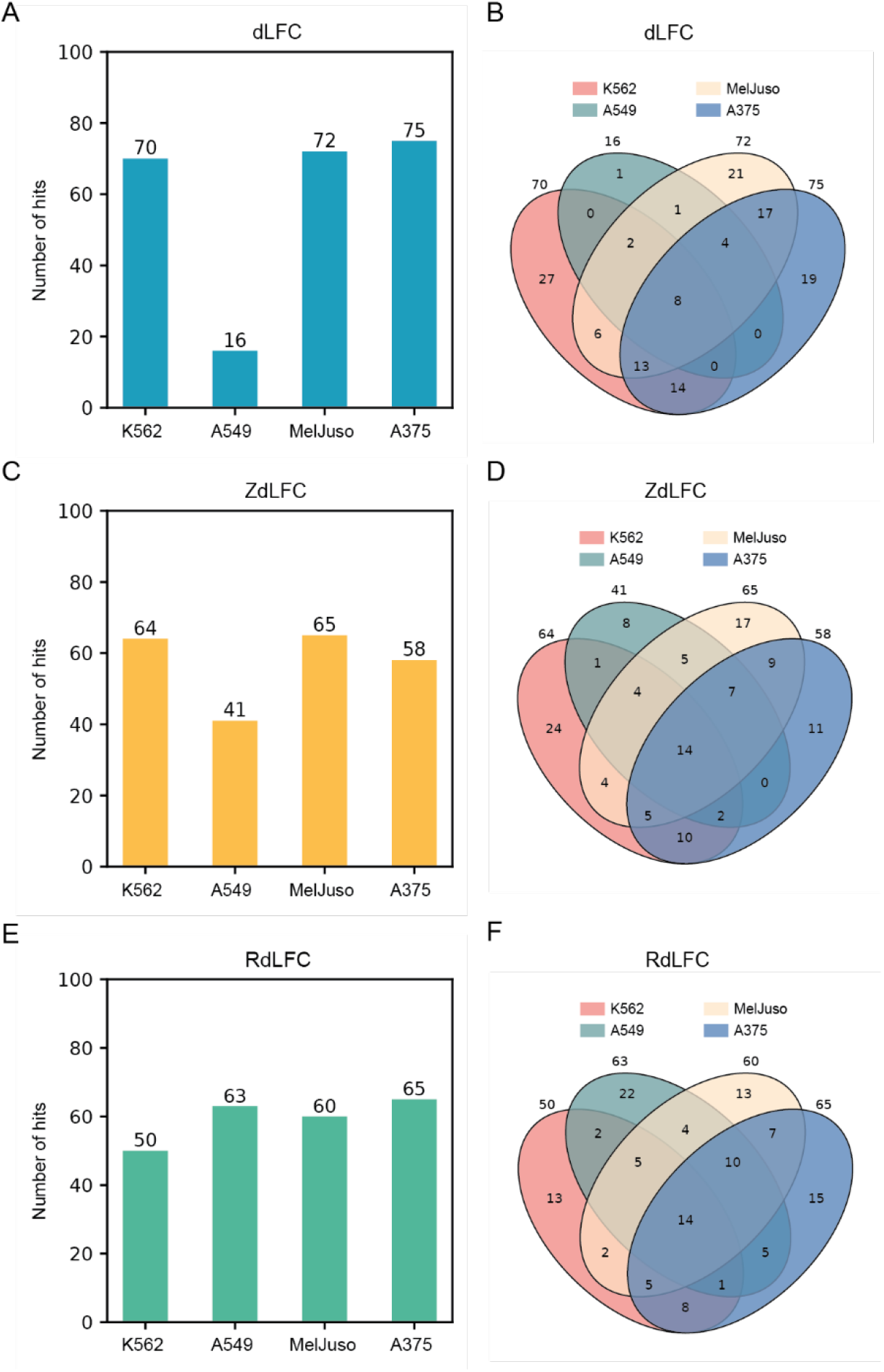
(A) Bar plot showing the number of synthetic lethal hits in each screen identified using the dLFC method with a threshold of dLFC < −1 and LFC < −1. (C) Bar plot showing the number of synthetic lethal hits in each screen identified using the ZdLFC method with a threshold of ZdLFC < −2 and ZLFC < −2. (E) Bar plot showing the number of synthetic lethal hits in each screen identified using the RdLFC method with a threshold of RdLFC < −0.7 and ZLFC < −2. (B, D, F) Venn diagrams illustrating the overlap of synthetic lethal hits identified in each of the four cell lines using the dLFC, ZdLFC, and RdLFC methods, respectively.

To measure consistency of hits across cell lines, we calculated the Jaccard coefficient of all pairs of screens (Figure 3A). With the thresholds we chose, the median Jaccard threshold for the ZdLFC approach is roughly equal to that of RdLFC, with both showing more consistency than the unscaled dLFC. We note that the ZLFC < −2 threshold for pairwise knockout essentiality appears to maximize Jaccard similarity across the data sets (Supplementary Figure 2A). Pairwise similarity of screens translates into groupwise similarity as well, as the number of hits in multiple screens is consistently higher with the Z-score and rescaling approaches (Figure 3B). The median pairwise sequence identity of synthetic lethals increases with the frequency of hits across cell lines (Figure 3C), although the variation is low. However, the distribution of pairwise sequence identity of hits in two or three out of four screens is almost identical to that of hits in four out of four screens (Supplementary Figure 2B). The Cohen’s D values are 0.06 and 0.03 for ZdLFC and RdLFC, respectively (hits in 3 vs 4 cell lines), and 0.17 and 0.10 for 2 vs 4 cell lines, with these very small effect sizes suggesting that sequence similarity might not be a good differentiator of context-dependent vs. pan-essential synthetic lethality. In contrast, hits with high pairwise sequence identity but observed in a single cell line contain clear examples of background-specific paralog synthetic lethals. For example, *NRAS/KRAS* are synthetic lethal in RTK-dependent cell line K562 while *KRAS* is singly essential in *KRAS* G12S mutant A549 cells (Figure 3E), and *MAPK1/MAPK3* are synthetic lethal in A549 and MelJuso, while K562 and A375 show specific dependence on *MAPK1*. Finally, the *CDK4/CDK6* pair is strongly synthetic lethal in MelJuso, though the gene pair was not tested in K562 and A549. A full list of paralog scores for each method is available in Supplementary Table 1.

**Figure 3.**
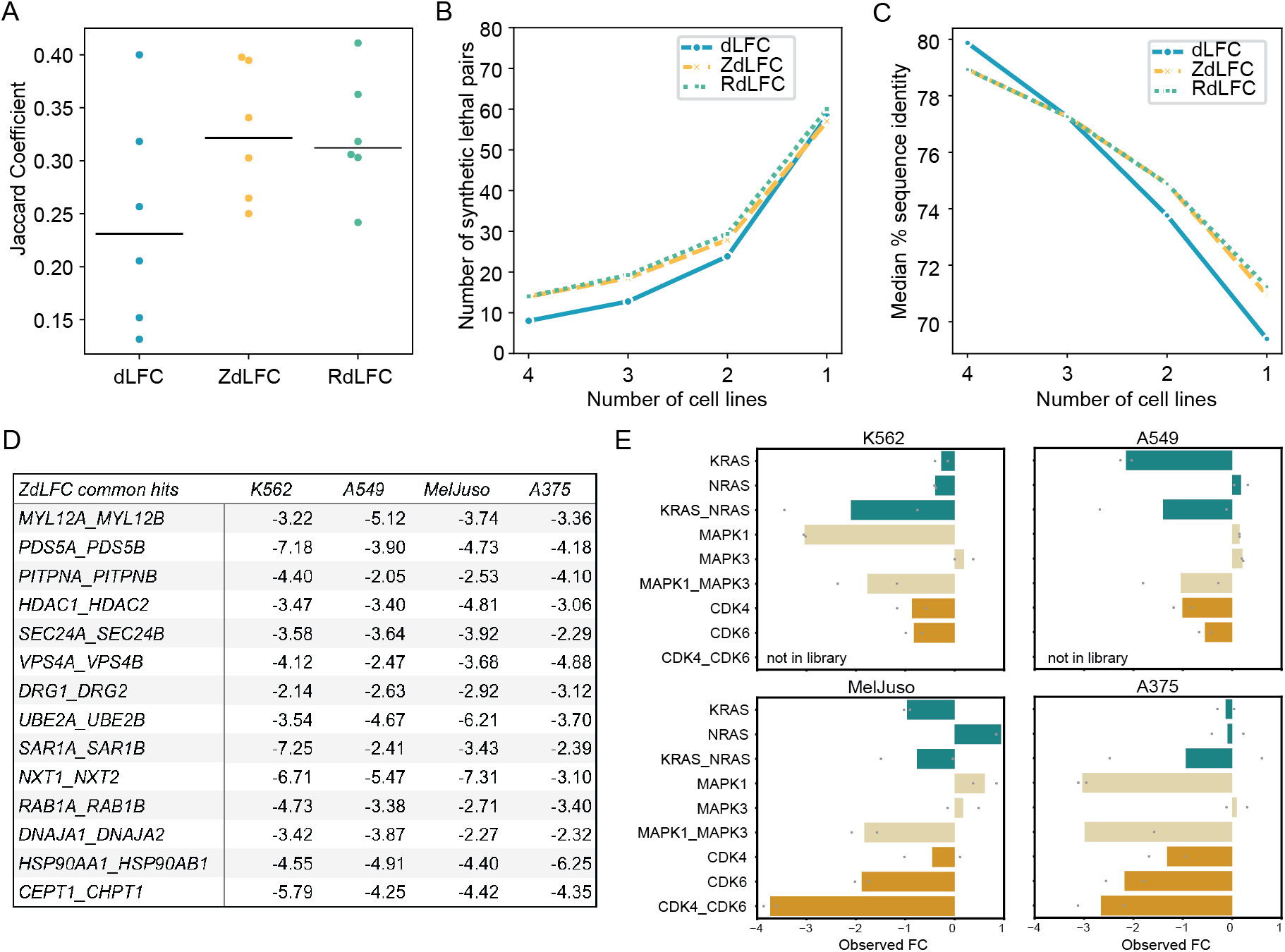
(A) Jaccard coefficients comparing the hits across all pairs of cell lines using three different methods. Black line indicates the median Jaccard coefficient for each method. (B) Line plot showing the mean number of synthetic lethal pairs identified in all four, three, two, and one screens using the three methods. (C) Line plot showing the median percent sequence identity of all synthetic lethal pairs identified in all four, three, two, and one screens using the three methods. (D) Table displaying the ZdLFC score of common hits identified across all four screens. (E) Background-specific paralog synthetic lethals shown in all four cell lines. Gene pair *CDK4/CDK6* was not included in the prototype library.

Given the above observations, we concluded that ZdLFC provides the best framework for scoring synthetic lethal paralogs. It provides more consistent hits across cell lines than the raw dLFC approach, in part because it can normalize for screens that show weaker overall distributions of fold change (e.g. A549 cells; Figure 2A and Supplementary Figure 1A). It yields virtually identical performance as the RdLFC (Figure 3A-C), without requiring a training set of known positive interactors – a requirement that must be met during experimental design.

We noted that the observed Jaccard coefficients for in4mer screens were significantly lower than those reported in Esmaeili Anvar et al^12^ for Cas12a-based paralog screens. We re-evaluated the Cas12a-based screen in Dede et al^6^ and the two recent SpCas9-based paralog screens^7,8^ using the ZdLFC approach and found generally similar performance as previously reported, though Parrish et al^7^ shows a marked improvement compared to our approach in Esmaeili Anvar et al due to the screen-specific data normalization (Figure 4). It is worth noting that a Jaccard coefficient of 0.33 corresponds to an intersection encompassing 50% of each of two equal-sized sets, and therefore indicates fairly strong coherence. Interestingly, the coherence of the Cas12a dual gRNA screens in Dede et al.^6^ remains substantially above that of the other screens, while the Cas12a in4mer screens look very similar to the SpCas9 dual gRNA screens. The Dede et al.^6^ test set was only 400 paralog pairs, while both Parrish et al.^7^ and Thompson et al.^8^ tested over 1,000 gene pairs, suggesting the high coherence of the Dede et al^6^ data was strongly influenced by the selection of the paralogs to assay. Conversely, the in4mer screen re-analysis here encompasses nearly twice as many target pairs (n=1,944) and uses about fivefold fewer reagents per gene pair, and results in roughly the same level of coherence as the Cas9 screens.

**Figure 4.**
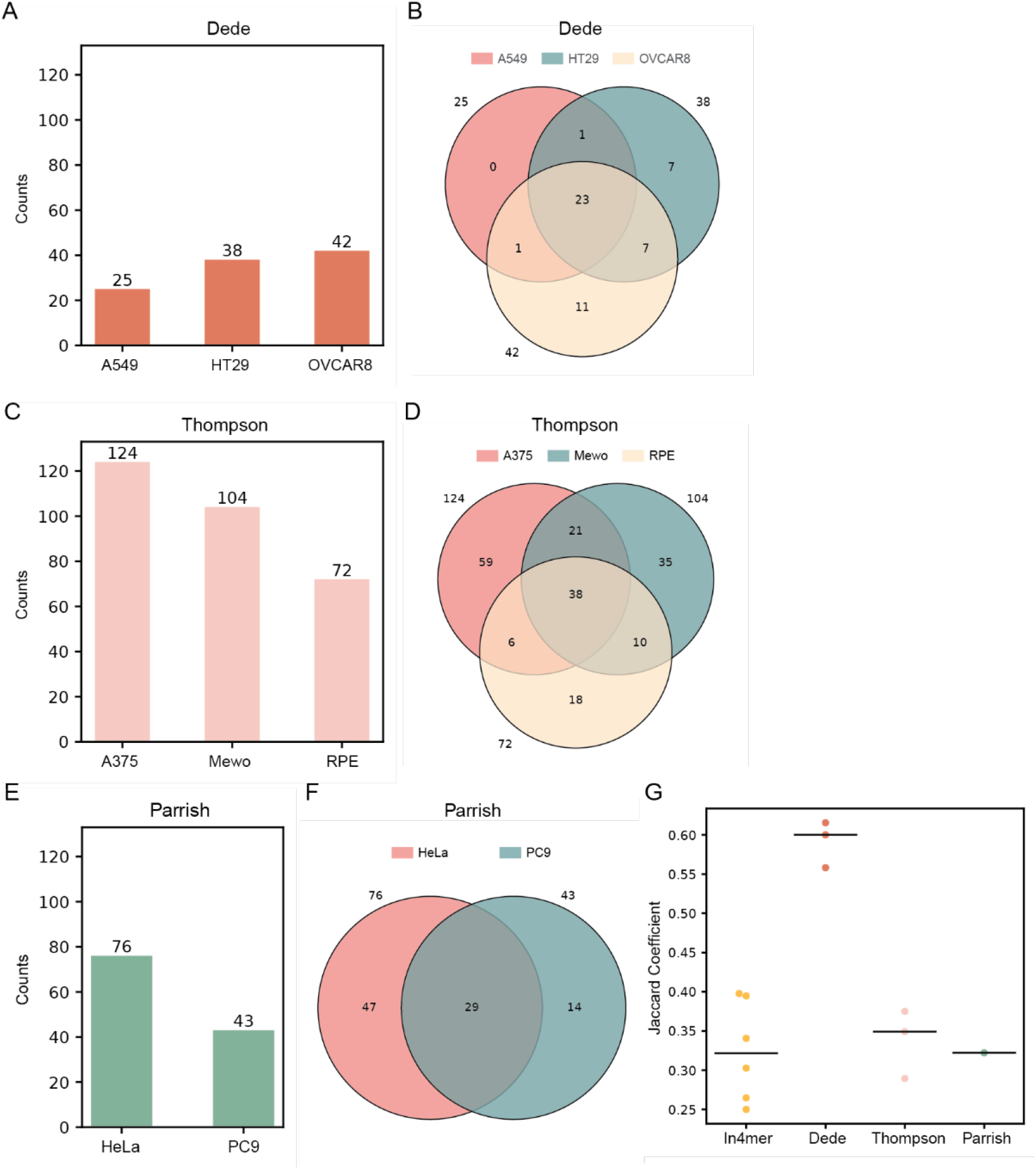
(A, C, E) Bar plots showing the number of synthetic lethal hits in each cell line from three other studies: a Cas12a-based screen^6^ and two SpCas9-based screens^7,8^. Hits were identified using the ZdLFC method with a threshold of ZdLFC < −2 and ZLFC < −2. (B, D, F) Venn diagrams illustrating the overlap of synthetic lethal hits identified in each cell line from the three studies using the ZdLFC method. (G) Jaccard coefficients comparing the hits across all pairs of cell lines within each study. Black line indicates the median of Jaccard coefficients for each study.

Finally, we provide a summary of hits across the screening platforms considered here. Supplementary Table 2 contains an unweighted count of the number of screens in which each synthetic lethal is observed. We reasoned that the near-equal Jaccard coefficients of the in4mer^12^, Parrish^7^, and Thompson^8^ platforms (Figure 4) and the apparent selection bias of Dede et al^6^ obviate the need for a weighted score as described in Esmaeili Anvar et al.

## Conclusions

Paralog synthetic lethality is of very high interest to the research community for several reasons. First, functional buffering by paralogs renders these gene families invisible to single-gene CRISPR knockout screens, revealing a knowledge gap in these large-scale efforts. Second, targeted therapies often inhibit related members of the same gene family, and in some cases rely on this multiple inhibition for efficacy (e.g. MEK, ERK inhibitors), thus making multiplex genetic inhibition a requirement for modeling drug efficacy. Third, and perhaps most importantly, functional buffering and polypharmacology extend beyond paralogs, but the constrained search space and relative frequency of paralog synthetic lethals make this an ideal testing area for more generalized genetic interaction technologies, both experimental and informatic.

To this end, we explored several bioinformatic options for analyzing paralog synthetic lethality data. We confirmed the importance of normalizing each data set, as has long been the case for single-gene CRISPR screens. Surprisingly, we find no benefit in using a training set of positive control synthetic lethals, although this may be simply because the training set is either too small or too noisy to be useful in this context. Overall, we find that ∼50% overlap between paralog synthetic lethal screens is a reasonable expectation for good quality screens, depending on the composition of the set of genes being analyzed.

Interestingly, this last point suggests that both the Cas12a and the Cas9 genetic interaction platforms are robust. Both implementations of the SpCas9 dual-promoter, dual-guide expression system gave largely equivalent results to the single-promoter, four-guide enCas12a system. We did not evaluate the multi-Cas systems as they add another layer of complexity that, in our view, is not justified, given the capability of the other platforms.

## Methods

### Prototype and Inzolia screens using the in4mer platform

The prototype library consists of 43,972 arrays targeting 19,687 single genes, 2082 paralog pairs, 167 paralog triples, and 48 paralog quads. The screenings were conducted on two cancer cell lines: K562 and A549. The Inzolia library consists of 50,085 arrays targeting 19,687 single genes, 4435 paralog pairs, 376 paralog triples, and 100 paralog quads. Screenings were performed on the MelJuso and A375 cell lines. More details of paralog selection and library construction can be found in the in4mer paper^12^.

The initial steps involved normalizing the raw read counts and assessing the overall quality of the screens. The read count data underwent preprocessing by adding a pseudo count of 5 reads to all arrays in each sample. The data was then normalized to a fixed total read count of 10 million reads. The guide-level log2 fold change (LFC) was calculated as the ratio of the normalized read count at the endpoint versus the T0 time point. Gene-level fold change (FC) was aggregated by averaging the guides. Both libraries included single knockout arrays targeting 50 essential genes as positive controls and 50 nonessential genes as the negative controls. These essential and nonessential genes were sourced from the Hart reference sets^15,16^. The quality of the screens was evaluated using Cohen’s D score, calculated as the mean FC difference between the essential and nonessential controls divided by the pooled standard deviation (Supplementary Figure 1A).

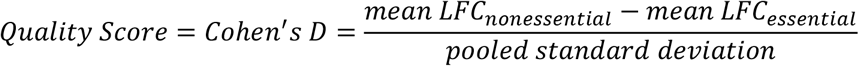

### Z transformed LFC

To calculate the z transformed LFC, the guide-level LFCs for each cell line were modeled using a two-components normal distribution with the GaussianMixture function from ‘sklearn.mixture’ in Python. The distribution with the higher weight represented the majority of guides that did not affect fitness, while the distribution with the lower weight, smaller mean, and larger variance represented a smaller number of genes whose knockout increased fitness defects. The mean and standard deviation of the higher-weight distribution were recorded and used to calculate the Z transformed LFC^17^. The equation used is as follows:

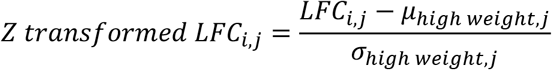

where *i* represents all arrays, including single genes and paralog pairs, j is the four cell lines, *µ*_*high weight,j*_ is the mean of the higher weight distribution, and *σ*_*high weight,j*_ is the standard deviation of the higher weight distribution (Supplementary Figure 1C).

### Three methods to score genetic interaction for paralogs

To identify the best method for detecting synthetic lethality in paralog screens, we analyzed the 1,944 common paralog pairs from the Prototype and Inzolia screens using three different quantification methods.

#### 1. Delta Log Fold Change (dLFC)

Genetic interactions were quantified by calculating the delta log fold change (dLFC), which is the log fold change of the pairwise gene knockout (observed) minus the sum of the single-gene knockout log fold changes (expected) (Figure 1A). This is referred to as the raw dLFC.

#### 2. Z-Transformed dLFC (ZdLFC)

To enhance accuracy, we calculated the Z-transformed dLFC (ZdLFC). We sought the best null model to fit the dLFC distribution by testing various components and determined that a single Gaussian component provided the best fit. However, we noticed the presence of outliers. To address this, we experimented with removing different quantiles and applied the standard outlier removal method: Q1 – 1.5*IQR, Q3 + 1.5*IQR. This method yielded the smallest mean distance between the empirical distribution function and the cumulative distribution function (CDF) fitted to our real data after outlier removal. Therefore, we used the fitted normal distribution, excluding outliers as defined by Q1 – 1.5*IQR, Q3 + 1.5*IQR as our null model.

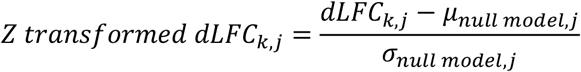

where *k* represents all common pairs. Q1 is the 0.25 quantile of dLFC distribution, Q3 is the 0.75 quantile of dLFC distribution, and IQR = Q3-Q1. *µ*_*null model,j*_ and *σ* _*null model,j*_ are the mean and standard deviation of the null model for cell line *j*. (Figure 1B; Supplementary Figure 1B)

#### 3. Rescaled dLFC (RdLFC)

The supervised RdLFC utilized the same null model as defined above. Additionally, it incorporated the observed dLFC of the 13 paralog synthetic lethal gold standards defined in Esmaeili Anvar et al.^12^ as a positive control reference set. We rescaled the dLFC of all pairs by setting the median of the positive control set to −1 and the mean of the null model to 0, adjusting the rest of the pairs accordingly.

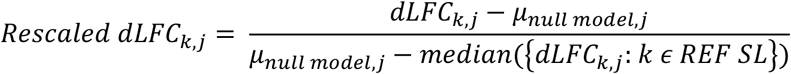

where *k* represents all common pairs, *j* represents the four cell lines, and REF SL represents the 13 paralog synthetic lethal gold standards.

### Evaluation of the three methods

To identify the most consistent method for calling synthetic lethal hits, we calculated the Jaccard similarity coefficient. We first generated all combinations of the four cell lines. Next, for each pair of cell lines, we calculated the fraction of the intersection of hits over the union of hits. The final Jaccard coefficient for each method was the median of all Jaccard coefficients from these pairwise comparisons.

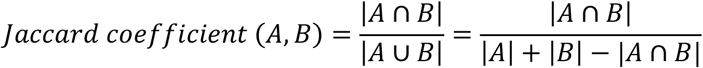

We also calculated the percent sequence identity of pairs identified as hits. Percent sequence identity data were obtained from BioMart. For each gene pair, the percent identity of paralogs AB and BA was recorded separately. We calculated the mean percent sequence identity for each gene pair (Figure 3C; Supplementary Figure 2B).

### Re-analysis of prior work

To re-evaluate the Cas12a-based screen in Dede et al.^6^ and the two SpCas9-based screens^7,8^ using the ZdLFC method, we followed these steps: first, raw read counts from the three studies were downloaded. The same preprocessing pipeline described above was applied to calculate gene-level LFC and dLFC. In Dede et al., the library targeted 403 paralog pairs and the screen was conducted in three cell lines: A549, HT29, and OVCAR8. The Thompson et al. study looked at 1,191 paralog and non-paralog pairs in the A375, Mewo, and RPE cell lines. Parrish et al. targeted 1,030 paralog pairs in HeLa and PC9 cell lines.

All LFC values in all screens were Z-transformed, and the dLFC values were Z-transformed as well. We used the same thresholds (ZdLFC<−2 and ZLFC<−2) to call hits in each study for fair comparison. To evaluate the consistency of hit calling across each method, we calculated the Jaccard coefficients (Figure 4G).

## Acknowledgements

JHC, NEA, and TH were supported by NIGMS grant R35GM130119 and NCI grant U01CA275886. TH is a CPRIT Scholar in Cancer Research and an Andrew Sabin Family Fellow. JC and JHC were additionally supported by R35CA274234. This work was supported by the National Cancer Institute’s Center for Cancer Genomics Cancer Target Discovery and Development (CTD^2^) initiative and by the NCI Cancer Center Support Grant P30CA16672.

---

**Supplementary Figure 1.**
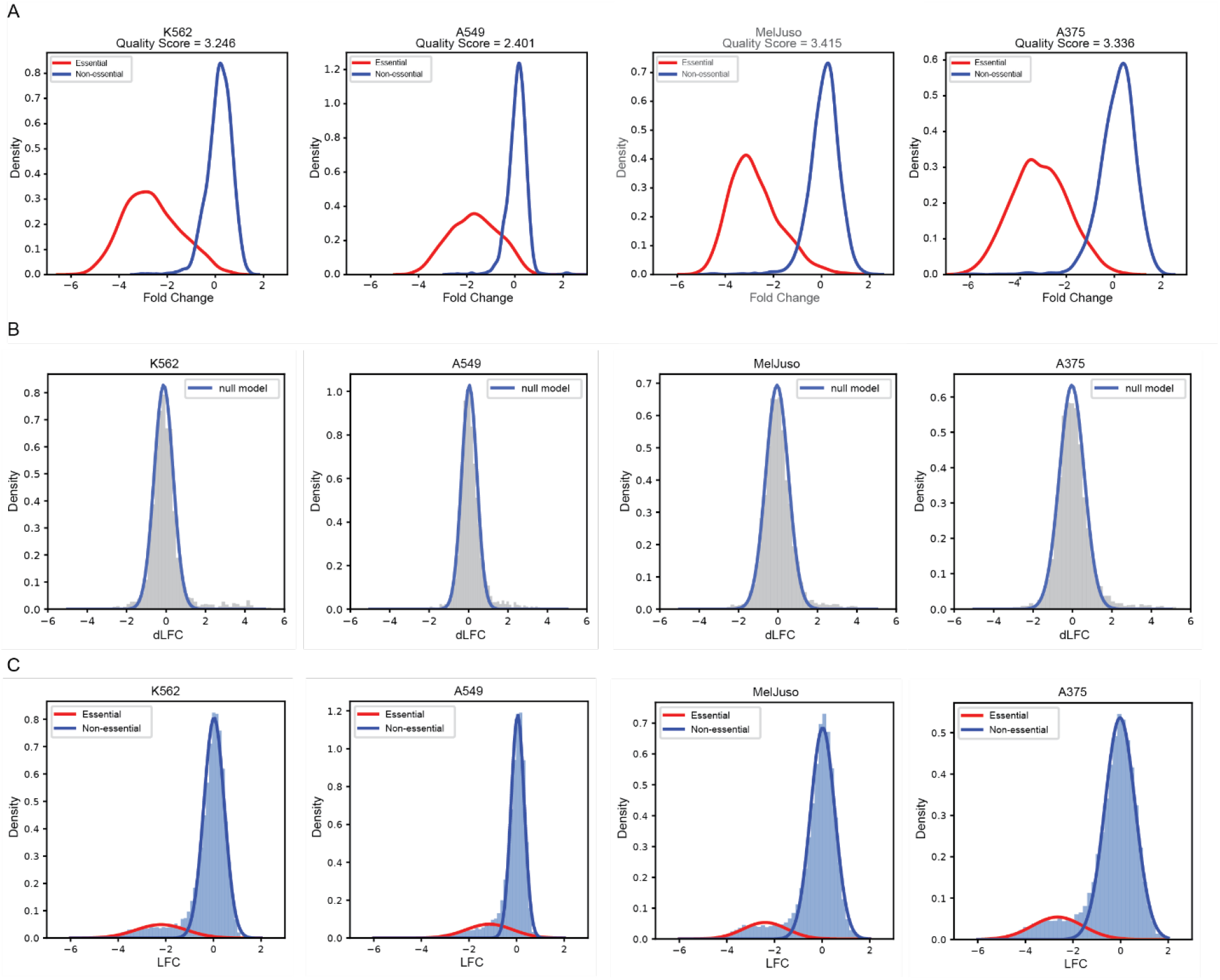
(A) Fold change distributions of arrays targeting reference essential (red) and non-essential (blue) genes, along with Cohen’s D quality score in four cell lines. This includes the prototype library in K562 and A549 cell lines, and the Inzolia library in MelJuso and A375 cell lines. (B) The dLFC histograms of the four cell lines with normal distribution fits after removing outliers (Methods). The blue curves represent the fit of the null model. (C) LFC histograms of the four cell lines, with a two-component Gaussian Mixture model representing the distribution. The red component models the essential genes, while the blue component represents the majority of genes that do not show severe fitness defects.

**Supplementary Figure 2.**
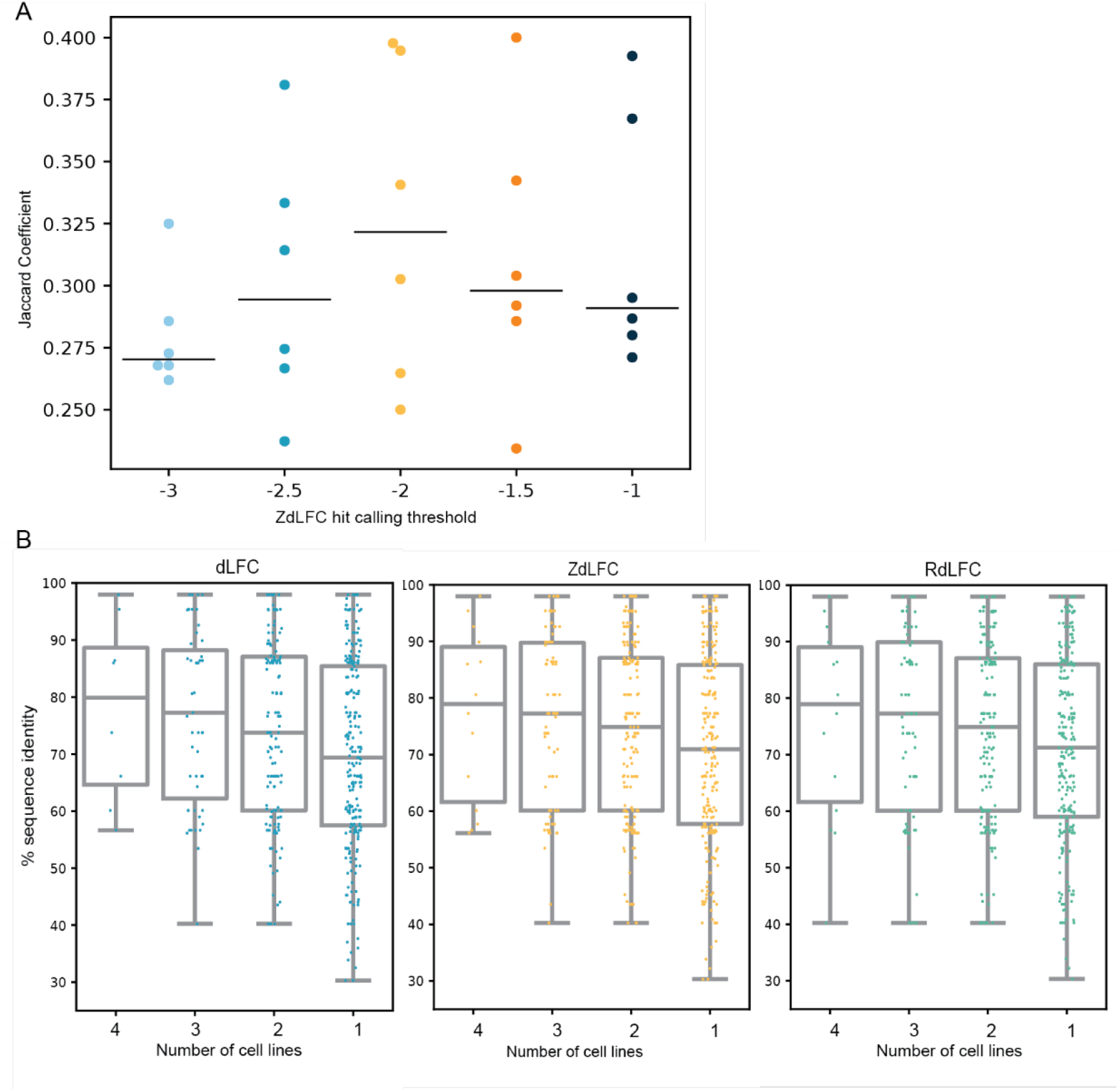
(A) Jaccard coefficients showing the hits across all pairs of cell lines with different thresholds using the ZdLFC method. Black line indicates the median Jaccard coefficient for each threshold. (B) Swarm plots combined with box plots showing the percent sequence identity of all synthetic lethal pairs identified in all four, three, two, and one screens using the three methods.

## References

1. Meyers, R. M. et al. Computational correction of copy number effect improves specificity of CRISPR–Cas9 essentiality screens in cancer cells. Nat Genet 49, 1779–1784 (2017).

2. Tsherniak, A. et al. Defining a Cancer Dependency Map. Cell 170, 564-576.e16 (2017).

3. Behan, F. M. et al. Prioritization of cancer therapeutic targets using CRISPR-Cas9 screens. Nature 568, 511–516 (2019).

4. Pacini, C. et al. A comprehensive clinically informed map of dependencies in cancer cells and framework for target prioritization. Cancer Cell 42, 301-316.e9 (2024).

5. De Kegel, B. & Ryan, C. J. Paralog buffering contributes to the variable essentiality of genes in cancer cell lines. PLoS Genet 15, e1008466 (2019).

6. Dede, M., McLaughlin, M., Kim, E. & Hart, T. Multiplex enCas12a screens detect functional buffering among paralogs otherwise masked in monogenic Cas9 knockout screens. Genome Biol 21, 262 (2020).

7. Parrish, P. C. R. et al. Discovery of synthetic lethal and tumor suppressor paralog pairs in the human genome. Cell Reports 36, 109597 (2021).

8. Thompson, N. A. et al. Combinatorial CRISPR screen identifies fitness effects of gene paralogues. Nat Commun 12, 1302 (2021).

9. Ito, T. et al. Paralog knockout profiling identifies DUSP4 and DUSP6 as a digenic dependence in MAPK pathway-driven cancers. Nat Genet 53, 1664–1672 (2021).

10. Gonatopoulos-Pournatzis, T. et al. Genetic interaction mapping and exon-resolution functional genomics with a hybrid Cas9–Cas12a platform. Nat Biotechnol 38, 638–648 (2020).

11. Zamanighomi, M. et al. GEMINI: a variational Bayesian approach to identify genetic interactions from combinatorial CRISPR screens. Genome Biol 20, 137 (2019).

12. Esmaeili Anvar, N. et al. Efficient gene knockout and genetic interaction screening using the in4mer CRISPR/Cas12a multiplex knockout platform. Nat Commun 15, 3577 (2024).

13. Hart, T. & Moffat, J. BAGEL: a computational framework for identifying essential genes from pooled library screens. BMC Bioinformatics 17, 164 (2016).

14. Kim, E. & Hart, T. Improved analysis of CRISPR fitness screens and reduced off-target effects with the BAGEL2 gene essentiality classifier. Genome Med 13, 2 (2021).

15. Hart, T., Brown, K. R., Sircoulomb, F., Rottapel, R. & Moffat, J. Measuring error rates in genomic perturbation screens: gold standards for human functional genomics. Mol Syst Biol 10, 733 (2014).

16. Hart, T. et al. High-Resolution CRISPR Screens Reveal Fitness Genes and Genotype-Specific Cancer Liabilities. Cell 163, 1515–26 (2015).

17. Lenoir, W. F. et al. Discovery of putative tumor suppressors from CRISPR screens reveals rewired lipid metabolism in acute myeloid leukemia cells. Nat Commun 12, 6506 (2021).

