## Supplementary Information for "Optimal methods for analyzing targeted pairwise knockout screens"

**Supplementary method**

Estimate genetic interactions with a Bayesian model – ParaBAGEL

ParaBAGEL is a supervised Bayesian model developed to estimate genetic interactions (GI) by utilizing delta log fold change (dLFC) metrics. is an extension of the BAGEL algorithm[1,2], updated to handle dual knockout multiplex CRISPR perturbation screens. As with BAGEL, the method employs positive and negative training sets. In this case, we employed the 13 paralog synthetic lethal (SL) gold standards from Esmaeili Anvar et al[3] as positive controls and empirically derived negative controls from the dLFC distribution. ParaBAGEL

Initial quality control procedures for gene-level LFC and dLFC were identical to those described in Methods. Positive controls (GI) were defined based on the 13-paralog synthetic lethal gold standards from Esmaeili Anvar et al. while negative controls (noGI) were derived from the null model described in Methods (Figure A).

The dLFC distributions of positive (GI) and negative (noGI) control gene pairs served as reference standards to label the deviation between having GI or not. Kernel density estimation was employed to model these distributions in the training sets. For each test pair, we calculated the likelihood that the observed dLFC originated from either the GI (red curve) or noGI (blue curve) training distributions (Figure A). Using a Bayesian supervised learning approach, we calculated a Bayes Factor (BF) score for each gene pair:

$$BF\ for\ GI = \frac{\Pr\,(D|k_{GI})}{\Pr\,(D|k_{noGI})} = \frac{\int \Pr\,(D|k\,,GI)\Pr(k|GI)\,dk}{\int \Pr(D|k,noGI)\Pr\,(k|noGI)dk}$$

where D represents the observed dLFC for a given gene pair and k denotes the dLFC distribution in the training set. A bootstrap resampling approach was used to randomly select gene pairs as the training sets. Considering the small sample size of the GI (n=13) control groups, we performed under-sampling for the no-GI group to balance the sample sizes. During each iteration, the k distributions were computed, and a BF for GI was calculated for each gene pair in the test set. The final score is the mean BF across all iterations.

The separation of GI and noGI distributions caused instability in the log ratio calculation (Figure B, green line). To mitigate this, we introduced overlapped normal distributions with smaller weights as pseudo-distributions on both the GI and noGI reference distributions. This adjustment aimed to enlarge the stable region for Bayes Factor calculation, optimizing four parameters: weights and standard deviations of the pseudo-distributions. The means of the pseudo-distributions were aligned with those of the GI and noGI distributions. Despite this, excessive reliance on pseudo-distributions compromised model performance.

Given the clear separation between GI and noGI distributions, simpler methods can effectively identify synthetic lethal pairs. Consequently, the Bayesian approach with pseudo-distributions was deemed unsuitable for this data set.

Supplementary method figures

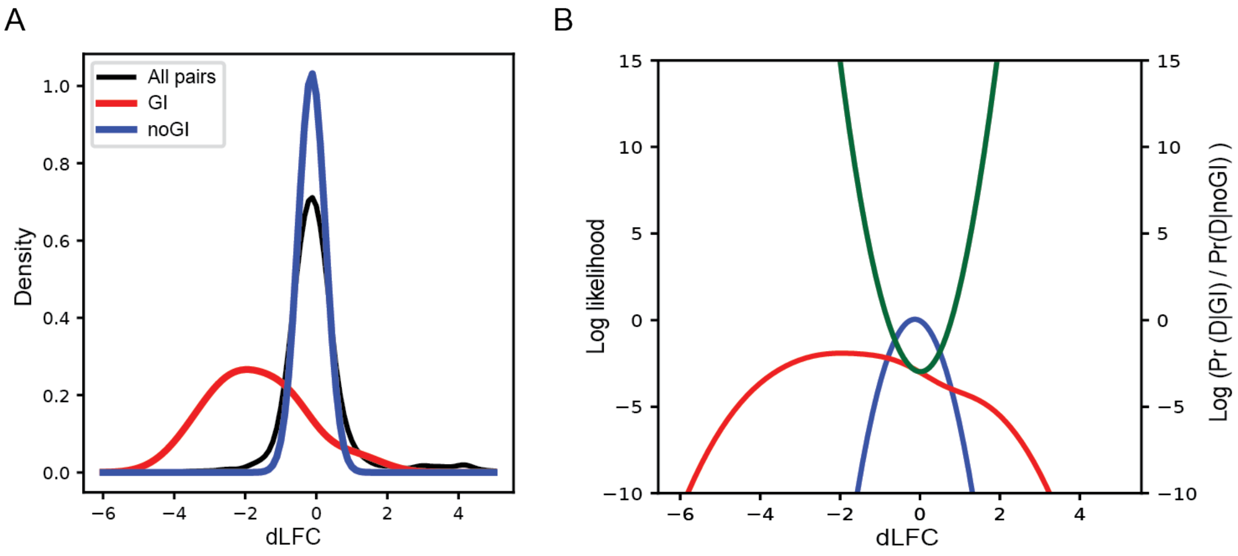


(A) The delta log fold change (dLFC) distributions from a single cell line. The red curve represents the kernel density plot of the 13-reference paralog synthetic lethal pairs (positive control), while the blue curve shows the kernel density plot of the null model (negative control). The dLFC values for all gene pairs are shown in black for reference. (B) The log likelihood functions for the red and blue curves from figure A are displayed on the left Y-axis. ParaBAGEL calculates the log likelihood ratio (green, right Y-axis) of these two curves. The green curve indicates that the stable region of the log ratio is narrow.

References


1. Hart, T. & Moffat, J. BAGEL: a computational framework for identifying essential genes from pooled library screens. *BMC Bioinformatics* **17**, 164 (2016).
2. Kim, E. & Hart, T. Improved analysis of CRISPR fitness screens and reduced off-target effects with the BAGEL2 gene essentiality classifier. *Genome Med* **13**, 2 (2021).
3. Esmaeili Anvar, N. *et al.* Efficient gene knockout and genetic interaction screening using the in4mer CRISPR/Cas12a multiplex knockout platform. *Nat Commun* **15**, 3577 (2024).
